# Assessing key decisions for transcriptomic data integration in biochemical networks

**DOI:** 10.1101/301945

**Authors:** Anne Richelle, Chintan Joshi, Nathan E. Lewis

**Affiliations:** Novo Nordisk Foundation Center for Biosustainability at the University of California, San Diego, School of Medicine, La Jolla, CA 92093, United States; Department of Pediatrics, University of California, San Diego, School of Medicine, La Jolla, CA 92093, United States; Department of Bioengineering, University of California, San Diego, La Jolla, CA 92093, United States

**Keywords:** omic data, systems biology, biochemical pathways, data integration

## Abstract

**Motivation:** To gain insights into complex biological processes, genome-scale data (e.g., RNA-Seq) are often overlaid on biochemical networks. However, many networks do not have a one-to-one relationship between genes and network edges, due to the existence of isozymes and protein complexes. Therefore, decisions must be made on how to overlay data onto networks. For example, for metabolic networks, these decisions include (1) how to integrate gene expression levels using gene-protein-reaction rules, (2) the approach used for selection of thresholds on expression data to consider the associated gene as “active”, and (3) the order in which these steps are imposed. However, the influence of these decisions has not been systematically tested.

**Results:** We compared 20 decision combinations using a transcriptomic dataset across 32 tissues and showed that definition of which reaction may be considered as active is mainly influenced by thresholding approach used. To determine the most appropriate decisions, we evaluated how these decisions impact the acquisition of tissue-specific active reaction lists that recapitulate organ-system tissue groups. These results will provide guidelines to improve data analyses with biochemical networks and facilitate the construction of context-specific metabolic models.

## Introduction

Most biological systems can be structured as networks, from cell signaling pathways to cell metabolism. These networks are invaluable for describing and understanding complex biological processes. For example, metabolic network reconstructions can illuminate the molecular basis of phenotypes exhibited by an organism, when used as a platform for analyzing data measuring gene expression, protein expression, enzymatic activity, or metabolite concentrations. For these analyses, the data are overlaid on the biological networks using Boolean rules that describe the relationship between the measured molecules (e.g., mRNAs, metabolites) and the network edges and nodes. These logical rules capture how the molecules influence each other’s activity (i.e., activation, inhibition, or cooperation), and allow users to quantify each network edge or define the status of each network component as either “on” or “off”. Therefore, a biological process can be described in a given context by adding or removing nodes and/or edges based on genome-scale data.

Genome-scale metabolic networks utilize this Boolean formulation connecting genes to reaction, and therefore have been used extensively as platforms for analyzing mRNA expression data to elucidate how changes in gene expression impacts cell phenotypes [1–8]. These studies have spanned diverse applications from identification of disease mechanisms [9,10] to identification of drug targets [11,12], and the evaluation of cell responses to drugs [13]. Despite the success of the many studies integrating omics data with biochemical networks, there are several challenges in the integration of omics data with networks that are infrequently discussed. These challenges impact the accuracy of context-specific networks, and include experimental and inherent biological noise, differences among experimental platforms, detection bias, and the unclear relationship between gene expression and reaction flux [14]. Furthermore, algorithmic assumptions influence the quality and functionality of resulting models and the physiological accuracy of their predictions [15–19].

While previous work has discussed the impact of various algorithms on obtaining physiologically accurate metabolic networks, the influence of the initial steps of data integration with biological networks has not been clearly evaluated and discussed in the literature. Thus, no universal rules have been established on how to integrate transcriptomic data, referred to here as “preprocessing”. These preprocessing steps include (1) how to account for network elements (e.g., reactions) that do not have a one-to-one relationship with genes and reactions (e.g., isozymes, complexes, and promiscuous enzymes), referred to here as *gene mapping*, and (2) how to define which genes are expressed or not, referred to here as *thresholding*, and (3) the order of gene mapping and thresholding in data integration. Here we evaluate the influence of the transcriptomic preprocessing steps and their consequences on the biological meaning captured by the data. Specifically, we do this by evaluating 20 different combinations of preprocessing steps, using transcriptomic data from 32 tissues. By evaluating the resulting 640 networks, we identify which decisions have the largest impact on network content, and which decisions best capture the similarities seen within tissues from the same organ-systems. This results in guidelines for overlaying transcriptomic data in metabolic networks and the lessons learned should be applicable to the analysis of transcriptomic data in all sorts of biological networks used for systems biology analysis.

## Results

### Preprocessing decisions for overlaying omics data on biochemical networks

Biochemical and other network types provide valuable platforms for analyzing and interpreting data. In these networks, links between nodes often represent enzyme-catalyzed reactions, and as such there is often not a one-to-one relationship between the genes and reactions. This relationship is represented using logical rules, referred as Gene-Protein-Reaction rules (GPRs, Supplementary Figure 1). When overlaying mRNA abundances on biochemical networks, GPRs are used to define which genes are the main determinants of the enzyme activity catalyzing a reaction. We refer here to this step as *gene mapping*. The most common assumption for multimeric enzyme complexes is that the gene with the minimum expression governs the activity. For isoenzymes, the activity may either depend on the total expression of all isoenzyme genes [20] or the isoenzyme gene with highest expression [21] (Figure 1A).

**Figure 1.**
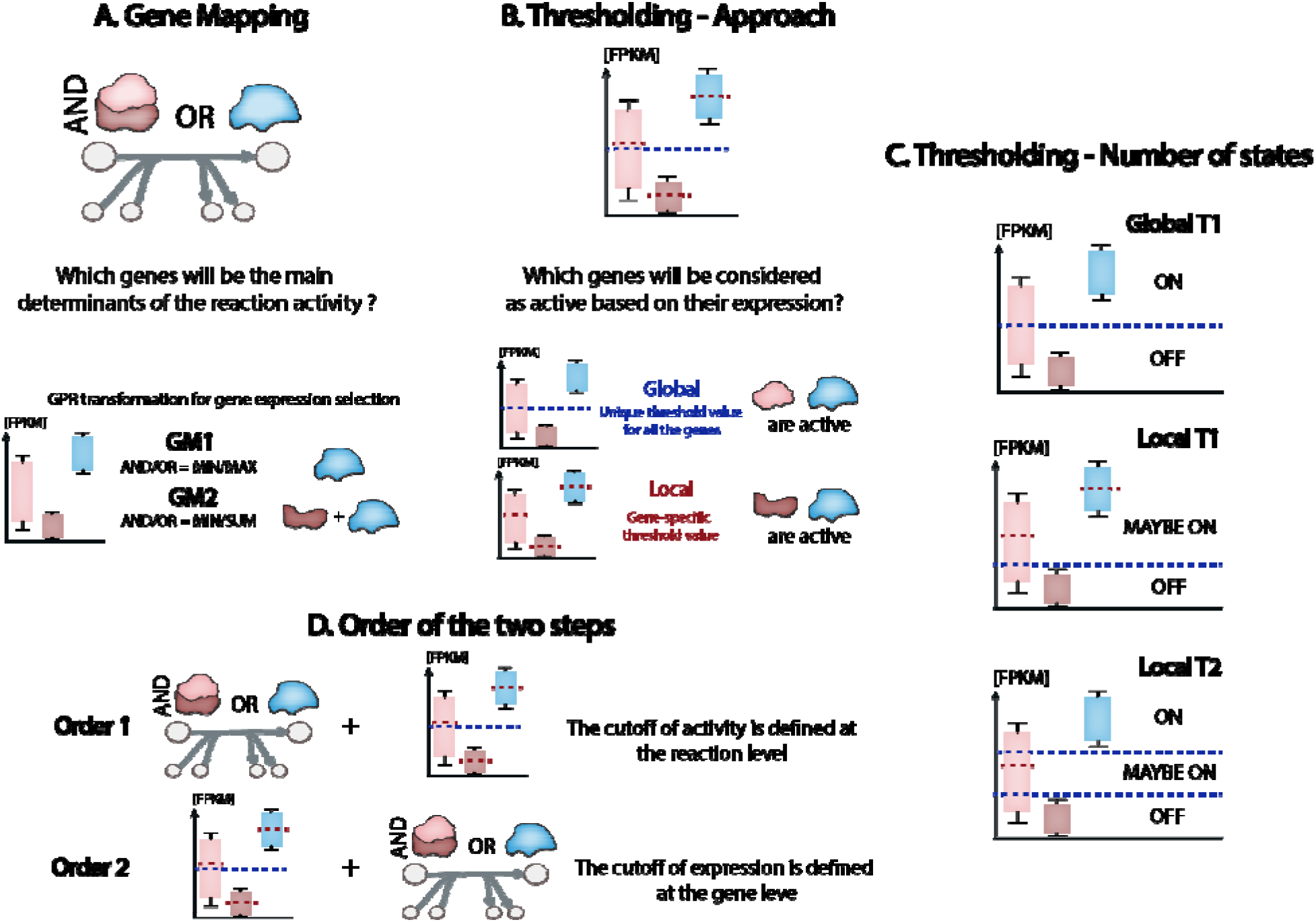
Formulation and implementation of preprocessing decisions. (A) Two types of gene mapping methods (GM1 and GM2) are compared. (B) Two types of thresholding approaches (global and local) are compared. (C) Formulation of three combinations of number of states (Global T1, Local T1, and Local T2) (D) Decisions about the order in which thresholding and gene mapping are performed. For Order 1, gene expression is converted to reactio activity followed by thresholding of reaction activity; for Order 2, thresholding of gene expression is followed by its conversion to reaction activity.

Furthermore, the absolute mRNA abundance is often considered to represent a gene’s potential activity by using a thresholding approach. That is, if the gene is expressed at a level above a threshold, it is often considered to be active. This threshold definition has been implemented in many different ways in the literature, from the use of only a single threshold to more complex rules involving multiple thresholds. For example, one unique threshold value can be applied to all genes (i.e., the global thresholding approach, [22,23]) while others have applied different thresholds to each gene (i.e., the local thresholding approach, [24,25]) (Figure 1B). When using one single threshold in a global context (i.e., global T1), the genes presenting an expression above this value are considered as active (i.e., ON) while the others are inactive (i.e. OFF) (Figure 1C). However, when multiple samples are available, one can compute a gene-specific threshold based on the distribution of the expression levels observed for this gene over all the samples (e.g., a local rule that sets a threshold equal to the mean expression level across all samples). In the literature, this gene-specific thresholding approach is often implemented in combination with a defined global threshold for genes presenting low expression values among all the samples (e.g., below the usual detection level associated to the measurement method) to prevent their inclusion with the active genes for some samples (i.e., local T1). Therefore, the genes whose expression is below the value defined by this global lower bound will always be considered as inactive (i.e., OFF), while other genes will fall under the local rule for gene-specific threshold definition (i.e., MAYBE ON) (Figure 1C). Another similar extreme case can be encountered when the gene expression level is high in all the samples. Therefore, we propose to also analyze the influence of using one lower and one upper threshold values defined based on the distribution of expression level off all the genes in all the samples (i.e. local T2). Doing so, the local rule for gene-specific threshold definition is actually applied only to the genes whose expression is between the range of values defined by the lower and upper bounds (i.e., MAYBE ON), ensuring that genes presenting low expression values among all the samples are always considered as inactive (i.e., OFF) while the ones with very high expression values among all the samples are always considered as active (i.e., ON) (Figure 1C).

Preprocessing of transcriptomic data for their integration into biochemical networks relies mainly on these two decisions: *gene mapping* and *thresholding*, but these can be implemented in different orders, with either gene mapping or thresholding occurring first (Figure 1D). Therefore, multiple combinations of these decisions could be made when overlaying data onto biochemical networks, and these decisions may influence the data integration and the subsequent biological interpretation (see Table 1, Figure 1, and a detailed explanation about the decisions presented in Methods section).

**Table 1:** Decisions involved in transcriptomic data preprocessing

| Decisions | Variables | Existing approaches | Biological meaning |
| --- | --- | --- | --- |
| **Gene Mapping** | GPR transformation for expression selection | AND/OR = MIN/MAX | Isoenzyme reaction activity is given by isoenzymes presenting the maximum activity |
|  |  | AND/OR = MIN/SUM | Isoenzyme reaction activity is given by the sum of the isoenzyme activities |
| **Thresholding** | Approach | local | Gene-specific threshold values |
|  |  | global | Unique threshold value for all the genes |
|  | Number of states | 2 states = 1 global threshold | OFF/ON |
|  |  | 2 states = 1 global threshold and 1 local rule | OFF/MAYBE ON |
|  |  | 3 states = 2 global thresholds and 1 local rule | OFF/MAYBE ON/ ON |
| **Order of the steps** | Gene Mapping (GM)<br>Thresholding (T) | GM + T | The cutoff of activity is defined at the reaction level |
|  |  | T + GM | The cutoff of expression is defined at the gene level |

Here, we integrated transcriptomic data from 32 different tissues in the Human Protein Atlas [25] with the Human genome scale model Recon 2.2 [26] using 20 different combinations of the 3 main preprocessing decisions (Table 1, Figure 1). This resulted in 640 different tissue-specific profiles of “expression” values for all gene-associated reactions in Recon 2.2. To specifically evaluate the immediate impact of the preprocessing decisions on the resulting networks, we focused our analysis on the content of the networks themselves (i.e., the definition of active biochemical pathways therein) and the biological interpretation of these networks.

### Active reaction sets are influenced by preprocessing decisions

Decisions regarding gene mapping, thresholding (i.e., approach and number of states), and order of steps affect the definition of active reaction sets. Specifically, the sets of active reactions (i.e., reactions with a non-zero expression level after overlaying the data) varied considerably in size from 358 reactions to 3286 reactions across all tissues, depending on preprocessing decisions and tissue type (Figure 2A). To assess the impact of each decision, we conducted a principal component analysis (PCA) of the reaction sets considered as active, depending on the preprocessing decisions (i.e., a PCA on the matrix of all active reactions vs. all combinations of decisions and tissues; see Methods for details). The first principal component explains >35% of the overall variance in active reaction content (Figure 2B). The thresholding related parameters (global/local and T1/T2) provide the most significant contribution to the variation in the first principal component (38.5%), with the differences between the global and local approaches having the greatest impact (Figures 2C, 2F, and Supplementary Figure 2). The effect of thresholding impacted the networks more than the differences across tissues, which only explained 15.8% of the variation in the first principal component. Tissue specific effects did not dominate until the second principal component, where it explained 73% of the variation in the component. The order of the preprocessing steps only provides a small contribution to the explained variation in the first principal component (Figure 2C, 2D). Meanwhile, the type of gene mapping has the least influence on active reaction sets (Figure 2C, 2E). These results indicate that the identification of active reactions is most heavily affected by the thresholding approach (as defined in Figure 1B), followed by the state definition used for thresholding (as defined in Figure 1C) and the order of preprocessing steps (as defined in Figure 1D) while the gene mapping method does not seem to have an influence.

**Figure 2.**
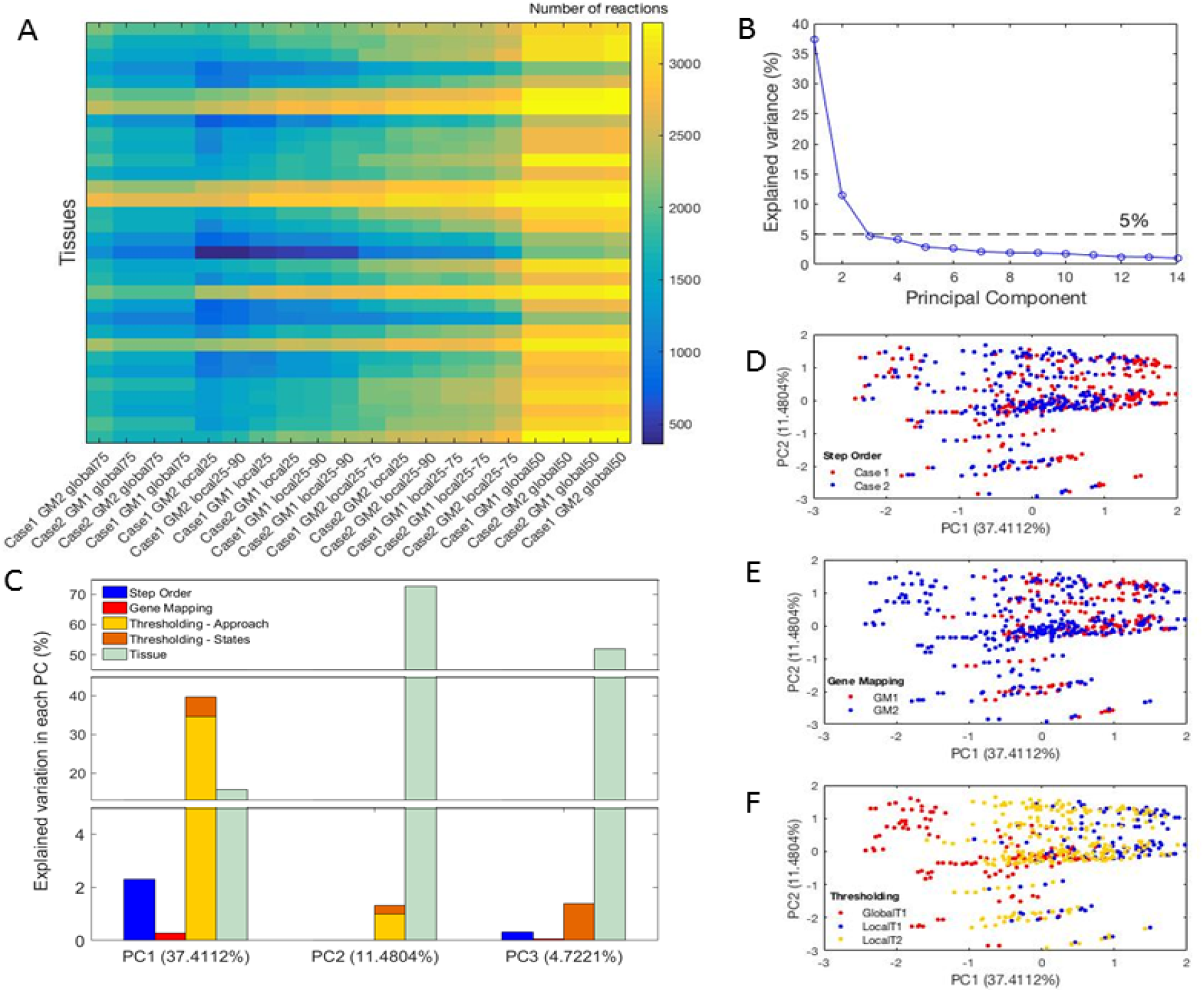
Preprocessing decisions affect the definition of active reactions sets. (A) Twenty different combinations of preprocessing decisions led to a large diversity number of reactions considered as active. (B) The first three principal components (PCs) explain most of the variance in the number of active reactions in a GEM. (C) Thresholding contributes the most to the first PC and more specifically the main contributor is the thresholding approach (i.e. local or global). (D, E and F). The influence of thresholding parameter selection is clear in the first PC (F), while the networks are less influenced by the gene mapping method (E) and the order of preprocessing steps used (D).

### Preprocessing decisions influence ability to capture tissue similarities within organ-systems

We assessed the similarities of tissues belonging to the same organ-system, based on the knowledge of the set of active reactions. We assumed that organ-system groups are formed by tissues working collaboratively to achieve a specific function (e.g., the gastrointestinal system turns food into energy). Therefore, we hypothesized that similarities of tissues within an organ system may lead to a more similar set of active metabolic reactions within the system, in comparison to other systems, as suggested by previous transcriptomic analyses [27,28]. To this end, we calculated Euclidean distances between pairs of tissues belonging to the same organ-system (Figure 3, Supplementary Figure 3, see Methods for more details). Our results highlight the influence of preprocessing decisions on the significance of tissue grouping at the reaction level. Moreover, we observed that some decisions improved the significance of tissue grouping: Order 2 works generally better than Order 1. Local T2 also is better than GlobalT1 and LocalT1. However, there was not a clearly superior approach for gene mapping in our analysis (Figure 4, Supplementary Figure 4).

**Figure 3.**
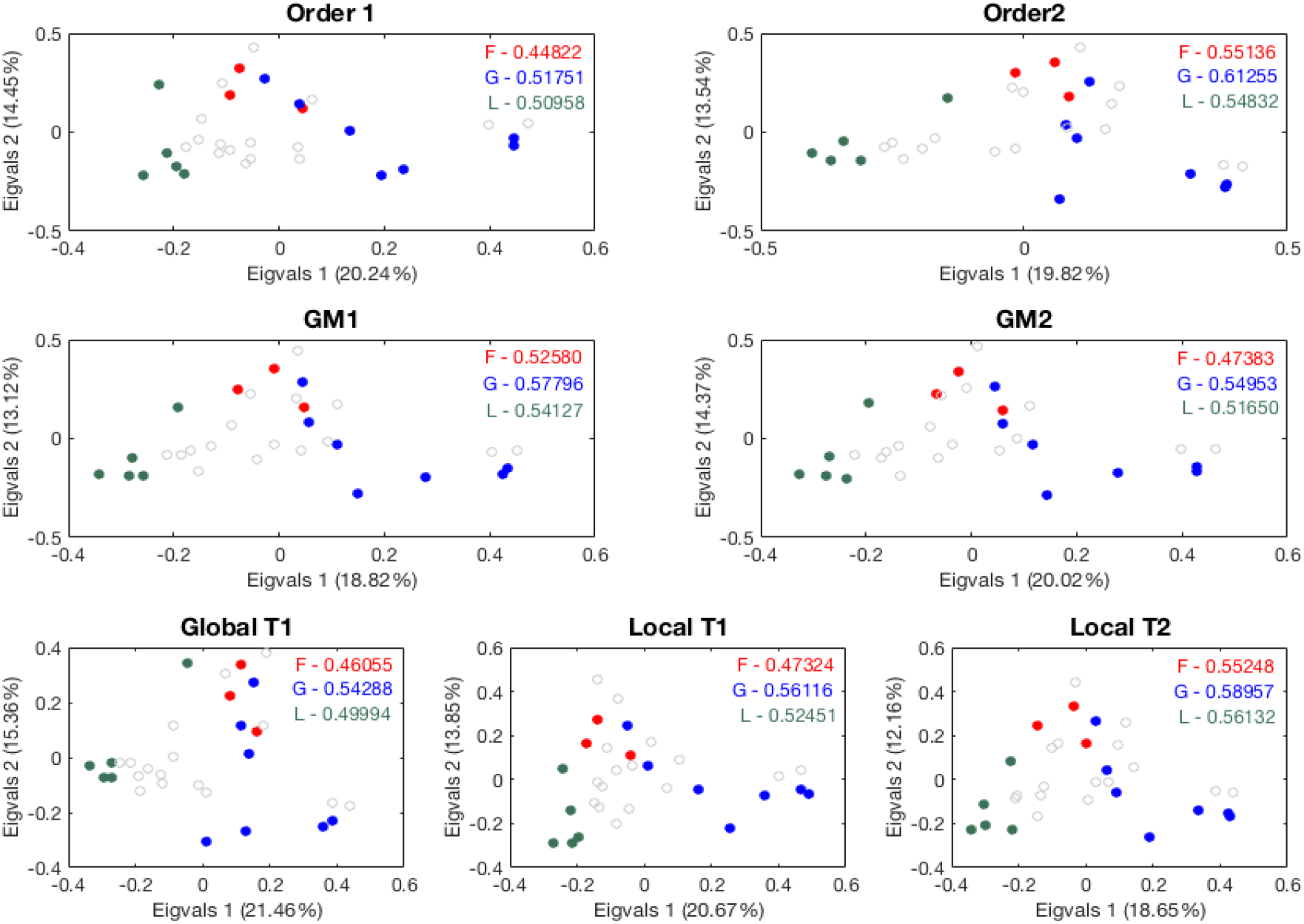
Influence of preprocessing decisions on capturing tissue similarities. Visual representation using a Principal Coordinates Analysis of the similarity between tissues grouped by organ system for each preprocessing decision (numbers in legends are the mean Euclidean distance of the tissues belonging to each group; F – Female reproductive group, G – Gastrointestinal group, and L – Lymphatic group).

**Figure 4.**
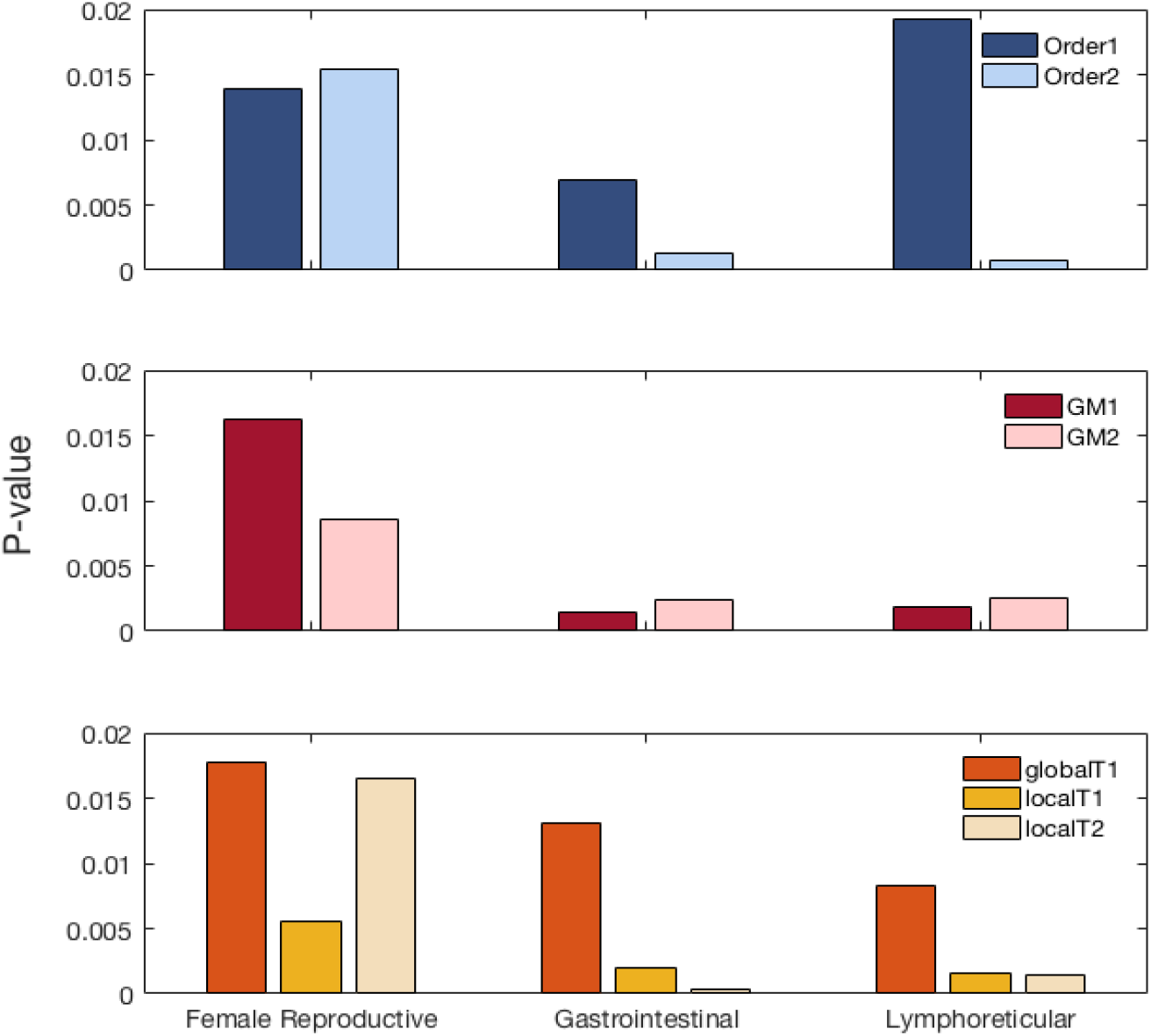
Preprocessing decisions influence the significance of tissue grouping at organ-system level. We compared the mean Euclidean distance observed between tissues belonging to the same organ-system to the mean Euclidean distance for 10000 randomly selected groups with the same number of tissues. The significance of the grouping (P-value) is computed as the proportion of random distances lower than the observed distance for each organ-system.

Some organ classification systems will group dissimilar organs together into a single organ-system, and we wondered if our analysis would still suggest the removal of such tissues from the organ-systems based on metabolic differences. For example, our previous analysis was done without associating the placenta to the *Female reproductive* organ-system group. However, the Human Protein Atlas groups it into the *Female reproductive* organ group (Supplementary Table 1). The placenta is functionally and histologically different from the other tissues of this group, being derived from both maternal and fetal tissue. This biological difference was successfully captured when we compared the tissue similarity analysis with and without the placenta in the *Female reproductive* organ-system group (Supplementary Figure 5).

## Discussion

Several methods have been developed to integrate transcriptomic data in GEMs, thus enabling the comprehensive study of metabolism for different cell types, tissue types, patients, or environmental conditions [8,12,22,23,29,30]. However, while these, and many other studies rely on preprocessing decisions to integrate the transcriptomic data in biochemical networks, each study makes different decisions without reporting the reason for their approach. Indeed, no rigorous comparison of the impacts of such decisions has been reported.

Here, we highlighted how different preprocessing decisions might influence information extracted from tissue specific gene expression data. We evaluated the influence of each preprocessing decision quantitatively by studying the active reaction sets and qualitatively by evaluating tissue grouping at an organ-system level. Our analysis suggested that thresholding related decisions have the strongest influence over the set of active pathways, and more specifically the thresholding approach (i.e., global or local; Figure 1C). This can be explained by the considerable influence of the decision on thresholding on the number of genes selected as expressed (Supplementary Figure 6). We note that threshold value choice for global thresholding was previously found to be the dominant factor influencing cell type-specific model content when context specific extraction methods were benchmarked [18]. When using global thresholds, the number of the genes selected to be active significantly decreases with increasing threshold value. However, the use of local thresholding leads to a smaller variation in the number of genes predicted to be active (Supplementary Figure 7). Furthermore, for similar state and value attribution (e.g. *T1 25^th^ percentile*), the use of the global thresholding approach leads to the selection of a larger number of genes predicted to be active in all tissues than the local approach (Supplementary Figure 6). Therefore, using a global threshold leads to fewer differences between tissues and a higher correlation of active reaction sets across tissues (Supplementary Figure 8), thus losing improved tissue specificity of the networks seen with the local thresholding approaches (Figure 4). This may have an important impact on analyses of tissue specific metabolism. Furthermore, the use of global thresholding is likely to lead to many false-negative reactions (i.e., reactions predicted to be inactive but are active), such as housekeeping genes that might be lowly expressed since they make essential vitamins, prosthetic groups, and micronutrients that are needed in low concentrations. Interestingly, the use of the T2 state definition seems to be less dependent on threshold values attributed than the T1 state definition when using a local approach (Supplementary Figure 7). Therefore, the use of a T2 state definition in combination of a local approach seems to successfully overcome the arbitrary aspect of threshold value selection and its influence on data preprocessing.

The order of preprocessing steps only moderately influences the definition of active reactions sets (Figure 1C). This decision implies two different interpretations of the influence of the RNA transcript levels on the determination of the enzyme abundance and activity associated to a given reaction. Indeed, the *Order 1* suggests that the measured expression levels determine the enzyme abundance available for a reaction while its associated activity will be defined depending on the gene chosen as the main determinant of the reaction behavior. On the other hand, the *Order 2* relies on a comparison of the activities of each gene associated with enzymes that might catalyze a reaction without directly accounting for the absolute transcript abundance. Our analyses suggest that *Order 2* provides more significant grouping for the *Gastrointestinal* and *Lymphoreticular* systems and does not considerably influence the grouping of the *Female reproductive* system. Advances in fluxomic measurement techniques will be invaluable to further investigate this preprocessing decision. Indeed, this would allow the analysis of the correlation between the RNA transcript levels and gene activity (expression data transformed using thresholding) of all the genes contributing to the definition of a reaction activity. Furthermore, this correlation analysis will further help with biological interpretation of this preprocessing decision and further refine guidelines for gene mapping decisions.

In our analysis, both gene mapping methods handle the AND relationships within a GPR rule in the same way but they differ in the treatment of OR relationships by either considering the maximum expression value (GM1) or a sum of expression values (GM2). Therefore, GM1 assumes that a reaction activity is determined by only one enzyme while GM2 accounts for the activity of all potential isoenzymes for a reaction. Surprisingly, while most of the reactions in Recon 2.2 are associated with at least two isoenzymes (Supplementary Figure 9A), the distributions of these reaction activities do not significantly change between the gene mapping approaches (Supplementary Figure 10). Indeed, even if there is a significant difference in the number of genes mapped to the model depending on the techniques used: an average of 58.3% of the genes present in the model and available in the HPA dataset are mapped to the model reactions using GM1 while 89.5 % are mapped using GM2. The expression value of genes that are unmapped using GM1 but mapped with GM2 is often below the 50^th^ percentile of the overall transcriptomic data available (Supplementary Figure 11) and therefore seems to not significantly influence the distribution of the reaction activities obtained. This is why the decisions relating to the gene mapping method do not influence the set of active reactions in the case of the transcriptomic dataset used in this study. However, it may not be the case for all transcriptomic datasets, especially if more metabolic genes are associated to high gene expression values. In this context, the development of more biologically meaningful gene mapping methods might be the key to capture differences between cell-types or tissues. Current gene mapping methods consider all enzymes as specialists (i.e. one enzyme is associated to one reaction). However, numerous enzymes are actually “generalists” as they exhibit promiscuity [31,32] (Supplementary Figure 9D). This functional promiscuity of an enzyme may be manifested in the form of competition between reactions catalyzed by this enzyme, and therefore influence the catalytic activity of an enzyme. In this context, future work may benefit from exploring strategies to handle enzyme promiscuity [33].

This benchmarking study emphasizes the importance of carefully evaluating the decisions and parameters for integrating transcriptomic data into biochemical networks. Indeed, numerous steps and decisions involved in the estimation of enzyme abundance and activity from transcriptomic data rely on biological assumptions that have not yet been leveraged. With the increasing availability and affordability of omic measurement techniques, studies filling the gap between mRNA expression and enzymatic activity will be of crucial importance.

## Conclusion

Decisions must be made on how to best handle and incorporate transcriptomic data into biochemical networks. Our benchmarking analysis of preprocessing decisions showed that thresholding approach influences the active reaction sets the most, even more than tissue-specific effects. Meanwhile, gene mapping has the lowest influence. We showed that some decisions better capture the functional tissue similarity across different organ systems. Overall, our analysis showed that transcriptomic data preprocessing decisions influence the ability to capture meaningful information about tissues. However, current preprocessing techniques present important limitations and decisions associated to this process should be made very carefully. In this context, development of more robust and biologically meaningful preprocessing techniques will be the key of the improvement of our understanding of tissue-specific behavior of an animal.

## Acknowledgements

This work was supported by generous funding from the Novo Nordisk Foundation provided to the Center for Biosustainability at the Technical University of Denmark (NNF16CC0021858), and from NIGMS (R35 GM119850). We also thank the W. M. Keck Foundation and the Lilly Innovation Fellowship program for generous funding that enabled this work.

## Methods

### Transcriptomic data

We used the Human Protein Atlas transcriptomic dataset (HPA) which includes RNA-Seq data of 20344 genes across 32 different human tissues [25]. Out of 20344 genes, 1663 can be mapped to the metabolic genes present in Recon 2.2 (99.4 % of coverage) [26]. Supplementary Table 2 presents the 10 genes of Recon 2.2 that are not associated with expression values in the HPA dataset and Supplementary Figure 12 presents the distribution of gene expression values in the HPA dataset.

### Genome-scale model of human metabolism – Recon2.2

Recon 2.2 [26] includes 1673 genes, 5324 metabolites and 7785 reactions. 3061 reactions do not have GPR associations. The remaining 4724 reactions are associated to 1797 different enzymes and about 20% of these reactions can be catalyzed by multiple isoenzymes. Almost 21% of the enzymes are formed by enzyme complexes (up to 46 subunits - reaction: NADH2_u10m) and about 54% of the enzymes are promiscuous enzymes (Supplementary Figure 9).

### Gene mapping

In metabolic networks, the relationship between genes and reactions is represented using logical rules, referred as Gene-Protein-Reaction rules (GPRs). These rules describe the association between the genes responsible for the expression of protein subunits forming the enzyme that catalyzes a reaction (AND for enzyme complexes; OR for isoenzymes). This relationship linking enzymes to reactions may have different types of GPR patterns. Some relationships are simple, with one gene encoding one enzyme that catalyzes one reaction. However, many are more complicated, in which one enzyme catalyzes multiple reactions (promiscuous), multiple proteins form an enzyme complex that catalyzes one reaction (multimeric), multiple enzymes catalyze one reaction (isoenzymatic), or multiple enzymes could catalyze multiple reactions (isoenzymatic promiscuous) [32] (Supplementary Figure 1). Gene mapping methods (GMMs) require combined use of the GPR rule and gene expression data to determine the enzyme activity associated to a reaction. In this regard, two methods have been used prominently in the field:

*i.* Selection of the *minimum* expression value among all the genes associated to an enzyme complex (AND rule) and the *maximum* expression value among all genes associated with an isoenzyme (OR rule). We refer to this method as GM1 [21].
*ii.* Selection of the *minimum* expression value among all the genes associated to an enzyme complex (AND rule) and *sum* of expression values of all the genes associated to an isoenzyme (OR rule). We refer to this method as GM2 [20].

### Thresholds

#### Thresholding Approaches

Thresholding approaches describe the scheme of threshold imposition on the gene expression value for a gene and/or reaction to be considered as “active”.

*i.* *Global approach*: The threshold value is the same for all the genes. The global approach is often applied when only one sample or condition is available and/or no information is available in the literature to define an expression threshold for a single gene. The “global threshold” is most often defined using the distribution of expression values for all the genes, and across all samples if multiple samples are available. This type of thresholding approach has been used, for example, in combination with a model extraction method called Gene Inactivity Moderated by Metabolism and Expression (GIMME) [22].
*ii.* *Local approach:* The threshold value is different for all the genes. The local approach is often applied when multiple samples are available as it allows a comparison of expression relative to many other samples and conditions. The “local threshold” for a gene is most often defined as the mean expression value of this gene across all the samples, tissues, or conditions [24,25,29].

The definition of thresholding criteria requires one to decide on how to partition the gene expression or reaction activity. In this regard, the ON/OFF state definition is often used in the literature. This type of state definition requires only one value to qualify if a gene/reaction is active. For example, when using one single threshold in a global context (i.e., hereafter referred as global T1), the genes presenting an expression above this value are considered as active (i.e., ON) while the others are inactive (i.e. OFF). However, this type of gene expression partition in a local context (e.g., expression threshold of a gene defined by its mean expression across all samples) presents limitations when facing genes with very low or very high expression values for all the samples. Indeed, when a gene presents always very low expression values, the use of the mean as threshold will lead to the consideration of its expression in some samples. Contrarily, some genes may be associated with very high expression values in all the samples. Doing so, while this gene should be considered as active, the current state partition will lead to considering this gene as non-expressed in all the samples presenting an expression value below the mean.

To overcome this problem, the gene-specific thresholding approach is often implemented in combination with the definition of a global threshold definition for genes presenting low expression values among all the samples (e.g., below the usual detection level associated to the measurement method) to prevent their definition as an active set for some samples (i.e., local T1). Therefore, the genes whose expression is below the value defined by this global lower bound will always be considered as inactive (i.e., OFF), while other genes will fall under the local rule for gene-specific threshold definition (i.e., MAYBE ON). The T1 state definition of local thresholding approach can be defined as follows “*the expression threshold for a gene is determined by the mean of expression values observed for that gene among all the tissues BUT the threshold must be higher or equal to a lower percentile bound globally defined”*.

Another similar extreme case can be encountered for genes with high expression values in all samples. To this end, an upper and a lower bound can be introduced to define the expression values for which a gene should always be considered as expressed or non-expressed. This will ensure that genes with very low expression values across all the samples will never be considered as active (i.e., OFF) and genes with very high expression across samples are always considered as active (i.e., ON). Doing so, the local rule for gene-specific threshold definition is applied only to the genes whose expression is in between the range of values defined by the lower and upper bounds (i.e., MAYBE ON). The definition of the local threshold with a T2 state definition can be expressed as follows: “*the expression threshold for a gene is determined by the mean of expression values observed for that gene among all the tissues BUT the threshold:(i) must be higher or equal to a lower percentile bound globally defined and (ii) must be lower or equal to an upper percentile bound globally defined.”*

#### Threshold values

The threshold values depend on the approach (i.e. local or global) and on the number of states (i.e. T1 or T2) used for thresholding. The global approach can only be associated with T1 state definition as it requires the assignment of only one threshold value. On the other hand, the local thresholding approach can be used in combination with either a T1 or a T2 state definition, as mentioned above. In the context of this study, we have chosen to compare the following combination of threshold value attribution:

*i.* *Global thresholding values:* The global threshold values chosen in this study are either the 50^th^ or the 75^th^ percentile (named respectively *Global T1 50^th^* and *Global T1 75^th^*). We also assessed the impact of using the 25^th^ and the 90^th^ percentiles. These thresholds exhibited tissue-specific sets of active genes that were more highly correlated and therefore decreased the ability to differentiate between tissues (*Global T1 25^th^*, Supplementary Figure 8) or more uncorrelated with more highly expressed genes removed, leading to a decreased ability to connect similar tissues (*Global T1 90^th^*, Supplementary Figure 6).
*ii.* *Local thresholding values:* we used the 25^th^ percentile of the overall gene expression distribution as lower bound for the local thresholding approach. This combination is referred as *Local T1 25^th^* when used alone. Note that, in the case of the HPA dataset, the 25^th^ percentile is equal to 1.2 FPKM and the detection limit of RNA-Seq technique is often considered at 1 FPKM (Supplementary Figure 12). Two different upper bounds have been used for the T2 state definition of the local approach: the 75^th^ (referred as *Local T2 25^th^ & 75^th^*) or the 90^th^ (referred as *Local T2 25^th^ & 90^th^*) percentiles of the overall gene expression distribution. The choice of the 75^th^ percentile as upper bound is based on the distribution of the mean expression value for each gene in the HPA dataset. Indeed, more than half of the genes are associated to a mean higher than the 50^th^ percentile of gene expression distribution (Supplementary Figure 13). To objectively assess the influence of the choice of this upper bound, we also compared the results obtained by imposing an upper bound of 90^th^ percentile.

### Ordering of preprocessing steps

The preprocessing for transcriptomic data could be done in two possible ways as described below:

*i.* *Order 1:* The gene expression values are associated to reactions using one of the gene mapping methods. These “reaction expressions” can further be used to define the set of active reactions by imposing thresholds of activity.
*ii.* *Order 2:* The thresholding is used to define the activity of each gene based on gene expression data. These “gene activities” are mapped to the reactions using one of the mapping methods. For Order 1, thresholding is imposed on these “reaction expressions” and no longer on the gene expression. This leads to the necessity to adapt the local threshold definition in the case of a preprocessing combination using Order 1 with GM2 gene mapping. Indeed, as the GM2 approach map multiple genes to a reaction, the activity of this reaction can no longer be defined by using the gene expression distribution. Therefore, the activity threshold for a reaction is determined by the sum of mean expression values observed for the genes mapped to this reactions) among all the tissues, but the mean expression value of each gene mapped to the reaction must be higher or equal to a lower percentile bound globally defined. Furthermore, it must be lower or equal to upper percentile bound globally defined.

### Principal Component Analysis (PCA)

A binary matrix is constructed in which each row represents one of the 20 preprocessing approaches and each column represents a variable: a reaction being active (1) or not active (0) in the GEM. The PCA analysis was conducted on this matrix after the removal of reactions being active for all or no preprocessing combinations and having each row centered to have zero mean.

### Assessment of tissues similarities

The set of active reactions have been used to compute the Euclidean distance between each tissue. We associated each tissue to an organ system using the classification proposed in the Human Protein Atlas (Supplementary Table 1) and computed the average Euclidean distance between tissues belonging to the same organ system. Note that, we only considered organ systems presenting more than two tissues within the same group (i.e. Female Reproductive, Lymphatic and Gastrointestinal). To compute the significance of our results, we generated the mean Euclidean distance for 10000 randomly selected group with the same number of tissues and computed the exact p-value (i.e. proportion of random distance lower than the observed distance) associated to each organ system.

### Data and software availability

The Human Protein Atlas transcriptomic dataset (HPA) were acquired from supplementary material of [25]. Recon 2.2 model was downloaded from https://github.com/u003f/recon2/tree/master/models. The matlab code for applying the different preprocessing combinations are available in COBRA toolbox v3.0, in the papers section [34].

